# Defining endogenous TACC3–chTOG–clathrin–GTSE1 interactions at the mitotic spindle using induced relocalization

**DOI:** 10.1101/2020.07.01.181818

**Authors:** Ellis L. Ryan, James Shelford, Teresa Massam-Wu, Richard Bayliss, Stephen J. Royle

## Abstract

A multiprotein complex containing TACC3, clathrin, and other proteins has been implicated in mitotic spindle stability. To disrupt this complex in an anti-cancer context, we need to understand the composition of the complex and the interactions between complex members and with microtubules. Induced relocalization of proteins in cells is a powerful way to analyze protein-protein interactions and additionally monitoring where and when these interactions occur. We used CRISPR/Cas9 gene-editing to add tandem FKBP-GFP tags to each complex member. The relocalization of endogenous tagged protein from the mitotic spindle to mitochondria and assessment of the effect on other proteins allowed us to establish that TACC3 and clathrin are core complex members and that chTOG and GTSE1 are ancillary to the complex, respectively binding to TACC3 and clathrin, but not each other. PIK3C2A, a membrane trafficking protein that binds clathrin, was previously proposed to also bind TACC3 and stabilize the TACC3–chTOG–clathrin–GTSE1 complex during mitosis. We show that PIK3C2A is not on the mitotic spindle and that knockout of this gene had no effect on the localization of the complex. We therefore conclude that PIK3C2A is not a member of the TACC3–chTOG–clathrin–GTSE1 complex. This work establishes that targeting the TACC3–clathrin interface or their microtubule-binding sites are the two strategies most likely to disrupt spindle stability mediated by this multiprotein complex.

## Introduction

During mitosis, chromosomes are segregated with high precision to generate two genetically identical daughter cells. This segregation is driven by the mitotic spindle, a bipolar microtubule array with associated motor and non-motor proteins (Manning and Compton, 2008). One non-motor protein complex that binds spindle microtubules contains TACC3, chTOG (also known as CKAP5) and clathrin (Fu et al., 2010; Hubner et al., 2010; Lin et al., 2010; Booth et al., 2011). This complex is important for stabilizing the bundles of microtubules that attach to kinetochores (kinetochore-fibers, k-fibers) by physically crosslinking them (Booth et al., 2011; Hepler et al., 1970; Nixon et al., 2015, 2017). Uncovering the molecular details of how proteins of this complex bind to one another and to microtubules is important to understand how i) mitotic spindles are stabilized and ii) we can target spindle stability in an anti-cancer context.

During mitosis, phosphorylation of TACC3 on serine 558 by Aurora-A kinase controls the interaction between clathrin and TACC3 (Booth et al., 2011; Cheeseman et al., 2011, 2013; Hood et al., 2013; Burgess et al., 2018). This interaction brings together the N-terminal domain of clathrin heavy chain and the TACC domain of TACC3 to make the microtubule-binding surface (Hood et al., 2013). Despite having a microtubule-lattice binding domain, chTOG is not needed for the complex to bind microtubules and interacts with the TACC3–clathrin complex via its TOG6 domain, binding to a stutter in the TACC domain of TACC3 (Booth et al., 2011; Hood et al., 2013; Gutiérrez-Caballero et al., 2015).

Despite this detail, the exact composition of the complex on kinetochore microtubules is uncertain. Besides TACC3, clathrin and chTOG, two further proteins, GTSE1 and Phosphatidylinositol 4-phosphate 3-kinase C2 domain-containing subunit alpha (PI3K-C2*α*/PIK3C2A) have been proposed to be members. Both were originally identified as binding partners for mitotic TACC3–clathrin (Hubner et al., 2010). Recent biochemical evidence convincingly shows that GTSE1 binds clathrin’s N-terminal domain and that this interaction localizes GTSE1 to spindle microtubules (Rondelet et al., 2020). Like chTOG, GTSE1 has the capacity to bind microtubules, but it appears to use TACC3–clathrin to bind the spindle (Monte et al., 2000; Scolz et al., 2012; Bendre et al., 2016). By contrast, PIK3C2A is a component of clathrin-coated vesicles where it acts as a lipid kinase (Gaidarov et al., 2001). It was recently proposed to act as a scaffolding protein in the TACC3–chTOG–clathrin complex by binding to both TACC3 and clathrin (Gulluni et al., 2017). PIK3C2A and GTSE1 bind to the same sites on clathrin’s N-terminal domain (Gaidarov et al., 2001; Rondelet et al., 2020) and although clathrin has the capacity to bind multiple proteins (Smith et al., 2017; Willox and Royle, 2012), this raises the question whether the binding of PIK3C2A and GTSE1 to TACC3–clathrin at the spindle is mutually exclusive. Dissecting this multiprotein complex is further complicated because each putative member is able to form subcomplexes that have different subcellular localizations (Gutiérrez-Caballero et al., 2015). TACC3–chTOG (without clathrin) localize to the plus-ends of microtubules (Nwagbara et al., 2014; Gutiérrez-Caballero et al., 2015). Similarly, GTSE1 binds plus-ends and can also stabilize astral microtubules of the mitotic spindle by inhibiting the microtubule depolymerase, MCAK/KIF2C (Scolz et al., 2012; Bendre et al., 2016; Tipton et al., 2017). PIK3C2A and clathrin are found in clathrin-coated vesicles away from the mitotic spindle (Gaidarov et al., 2001). Biochemical approaches do not have the capacity to discriminate these subcomplexes from the multi-protein complex on K-fibres. Therefore, subcellular investigation of protein interactions are required to answer this question.

Knocksideways is a method to acutely and inducibly relocalize a protein to mitochondria in order to inactivate that protein (Robinson et al., 2010). In the original method, the target protein is depleted by RNAi and an FKBP-tagged version is expressed alongside MitoTrap (an FRB domain targeted to mitochondria), relocalization is achieved by the addition of rapamycin. This method has many advantages over slow inactivation methods such as RNAi knockdown or gene disruption (knock-out) approaches (Royle, 2013). We have previously used knocksideways in mitotic cells to investigate protein-protein interactions since any proteins that are in a complex with the target protein also become mislocalized to the mitochondria (Cheeseman et al., 2013; Hood et al., 2013). This approach has the added advantage that the subcellular location of proteins can also be tracked during the experiment, and that it can be done at specific times; allowing us to pinpoint where and when interactions occur.

In this study, we applied a knocksideways approach to investigate how proteins of this complex bind to one another and to microtubules of the mitotic spindle. Instead of overexpression and RNAi, we sought to tag each target protein with FKBP and GFP at its endogenous locus using CRISPR/Cas9-mediated gene editing. This strategy allows us to study these subcellular interactions at the endogenous level for the first time. The cell lines we have created are a multi-purpose “toolkit” for studying microtubule-crosslinking proteins by live-cell imaging, biochemistry, or electron microscopy (Clarke and Royle, 2018).

## Results

### Generation and validation of clathrin, TACC3, chTOG and GTSE1 knock-in HeLa cell lines

Our first goal was to tag four proteins with FKBP and GFP at their endogenous loci using CRISPR/Cas gene editing. Clathrin (targeting clathrin light chain a, LCa/CLTA), TACC3, chTOG/CKAP5, and the clathrin-interacting protein, GTSE1 were edited in HeLa cells so that they had a GFP-FKBP tag at their N-terminus or an FKBP-GFP tag at their C-terminus (Figure 1A). The dual FKBP and GFP tag allows direct visualization of the protein as well as its spatial manipulation using knocksideways (Cheeseman et al., 2013; Robinson et al., 2010). Following editing, GFP-positive cells were isolated by FACS and were validated using a combination of PCR, sequencing, Western blotting (Figure 1B, Supplementary Figure S1A), and fluorescence microscopy (Figure 1C, Supplementary Figure S1B). These validation steps yielded a cell line for each protein that could be used for all future analyses. Homozygous knock-in was achieved for CLTA-FKBP-GFP, GFP-FKBP-TACC3, and GTSE1-FKBP-GFP. Despite multiple attempts to generate a homozygous knock-in for chTOG-FKBP-GFP, we only recovered heterozygous lines (more than twenty heterozygous clones in three separate attempts). We assume that homozygous knock-in of chTOG-FKBP-GFP is lethal.

**Figure 1.**
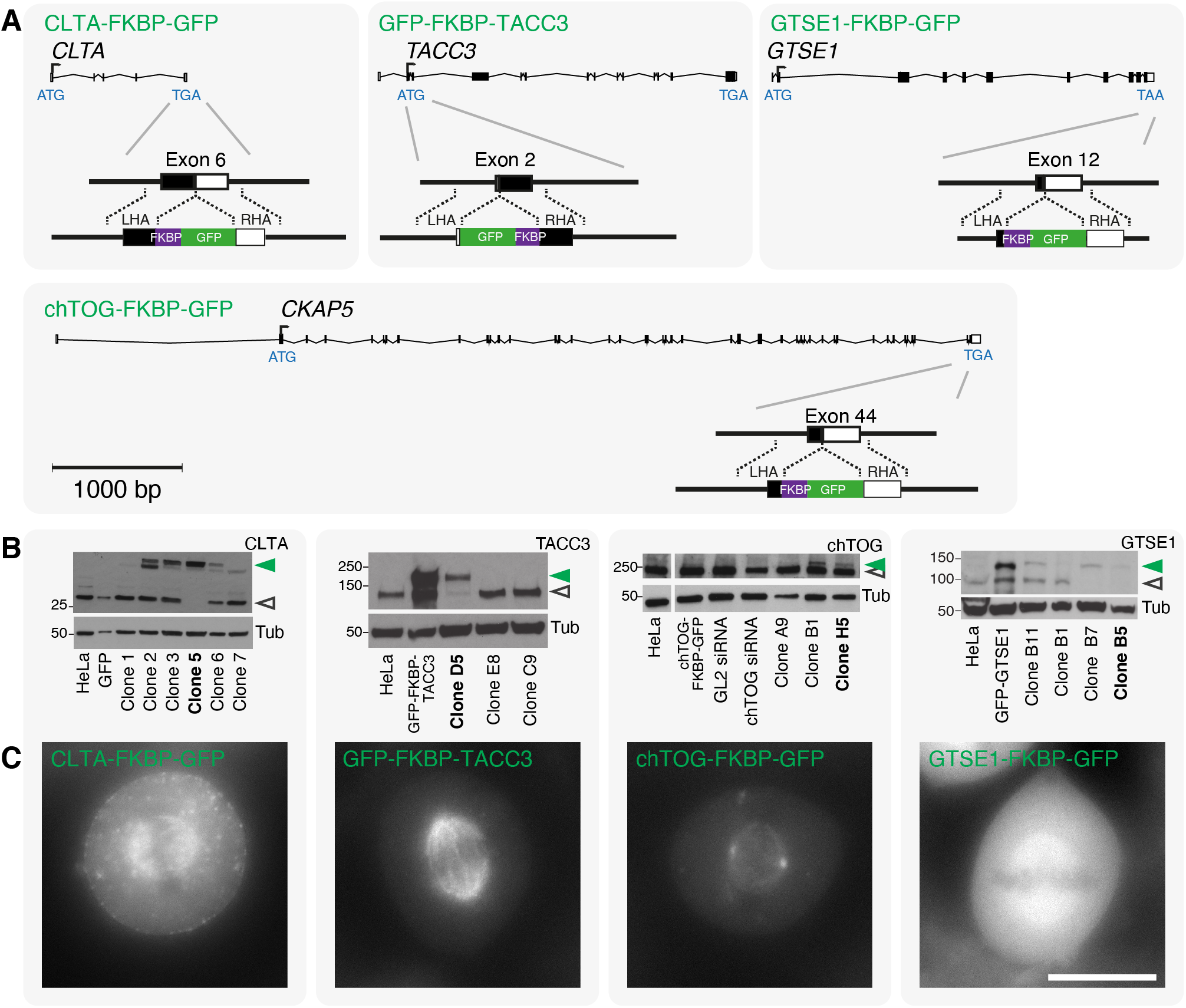
Generation of knock-in HeLa cell lines using gene editing. (**A**) Strategy to tag clathrin (CLTA), TACC3, chTOG/CKAP5 and GTSE1 with FKBP and GFP at their endogenous loci. Cas9n D10A nickase was used to target the indicated site and a repair template with FKBP-GFP or GFP-FKBP tag flanked by left and right homology arms (LHA/RHA) was used as indicated. Scale bar, 1000 bp. GFP-positive cells were individually sorted by FACS and validated using a combination of Western blotting (B), imaging (C) and DNA sequencing (not shown). (**B**) Western blotting showed negative clones and positive clones that were either homozygous (single band, shifted by ~30 kDa) or heterozygous (two bands, one at expected size and the other shifted by ~30 kDa) knock-in cell lines. The tagged and untagged proteins are denoted by filled green and open gray arrowheads, respectively. Clones used in this work are highlighted in bold. HeLa cells may have more than two alleles of the targeted gene. We use the term homozygous to indicate editing of all alleles and heterozygous to indicate that at least one allele was edited and that an unedited allele remained. PCR and DNA sequencing confirmed that: CLTA-FKBP-GFP (clone 5) is homozygous, GFP-FKBP-TACC3 (clone D5) is homozygous, chTOG-FKBP-GFP (clone H5) is heterozygous, and GTSE1-FKBP-GFP (clone B5) is homozygous, (**C**) Micrographs of each tagged cell line indicating the correct localization of each tagged protein. Scale bar, 10 μm.

The localization of tagged proteins in all cell lines was normal. In mitotic cells, clathrin is located on the spindle and the cytoplasm/coated pits, TACC3 is located exclusively on the spindle, chTOG is located on the spindle but more pronounced on the centrosomes and kinetochores, and GTSE1 is localized throughout the spindle and the cytoplasm (Figure 1C, Supplementary Figure S1B), consistent with previous observations (Gergely et al., 2000, 2003; Royle et al., 2005; Foraker et al., 2012; Bendre et al., 2016). Overexpression of TACC3 can result in the formation of aggregates (Gergely et al., 2000; Hood et al., 2013) which have recently been described as liquid-like phase-separated structures(So et al., 2019). We note that at endogenous levels in HeLa cells, GFP-FKBP-TACC3 does not form these structures (Figure 1C, Supplementary Figure S1C).

As a further validation step, we assessed mitotic timings in each knock-in cell line and found progression to be comparable to their respective parental HeLa cells. These observations indicate that the addition of an FKBP and GFP tag did not affect the mitotic function of clathrin, TACC3, chTOG, or GTSE1 and that clonal selection did not adversely affect mitosis in the four cell lines (Supplementary Figure S1D).

Finally, we assessed the functionality of the FKBP moiety by performing knocksideways experiments (Figure 2A). Each cell line, expressing mCherry-MitoTrap was imaged live during the application of 200 nM rapamycin. At metaphase, CLTA-FKBP-GFP, GFP-FKBP-TACC3, chTOG-FKBP-GFP, and GTSE1-FKBP-GFP were all removed from the spindle and relocalized to the mitochondria by rapamycin addition (Figure 2B). The time course of removal was variable but was complete by 10 min. (Supplementary Video SV1,SV2,SV3,SV4). In summary, the generation and validation of these four knock-in cell lines represents a toolkit that we can use to study clathrin, TACC3, chTOG and GTSE1 at endogenous levels (see Table 1).

**Figure 2.**
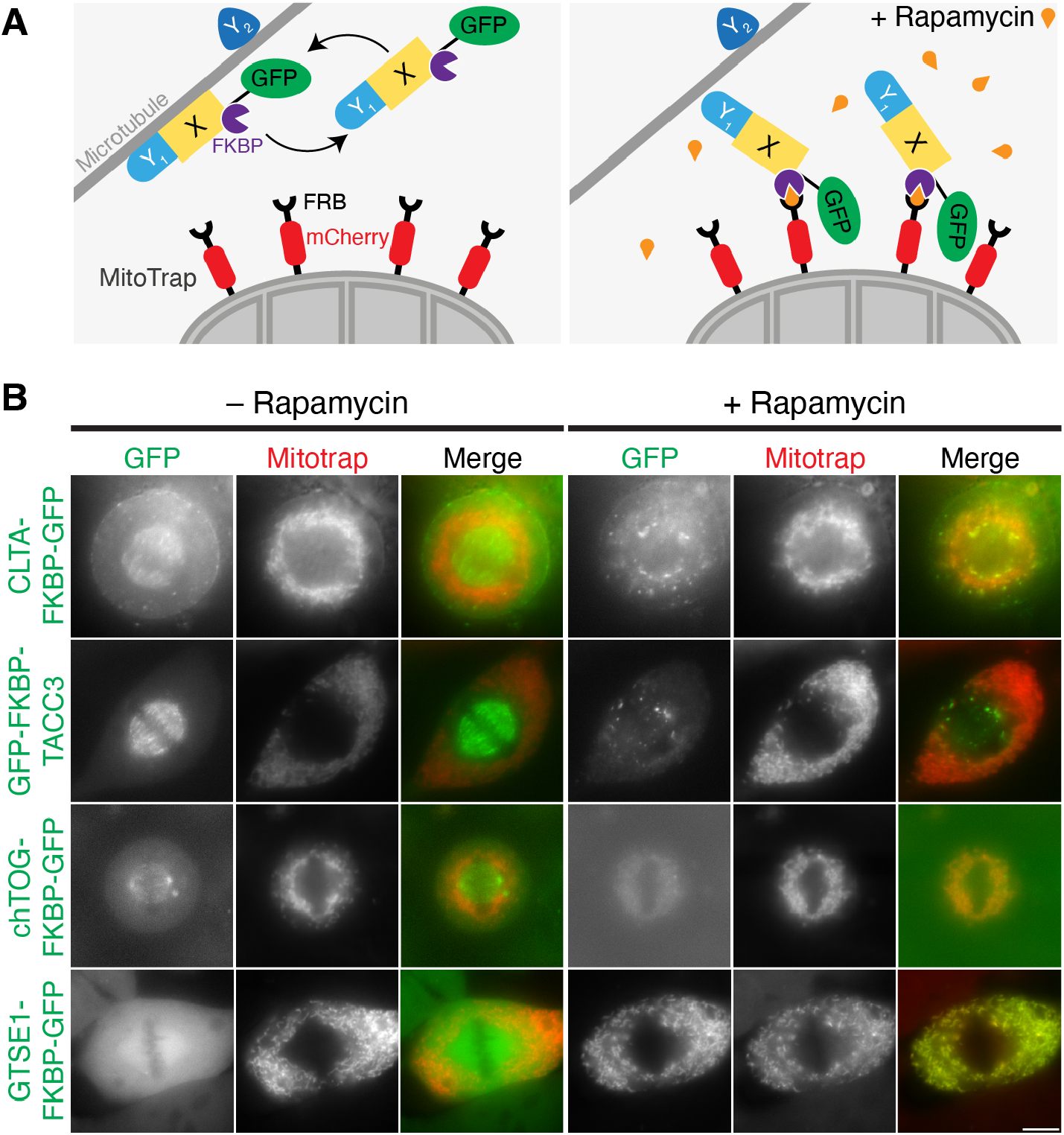
Generation of knock-in HeLa cell lines using gene editing. (**A**) Schematic diagram of knocksideways in gene edited cells. A microtubule-binding protein X is fused to FKBP and GFP. MitoTrap, an FRB domain targeted to mitochondria, tagged with mCherry, is transiently expressed. Addition of rapamycin causes the relocalization of proteins to the mitochondria (Robinson et al., 2010). This strategy can also be used to assess if another protein Y, co-reroutes with X to the mitochondria. *Y*_1_ co-reroutes with X, indicating that they form a complex, whereas *Y*_2_ does not. (**B**) Live cell imaging of knocksideways of gene edited cell lines. Indicated tagged cell lines expressing were imaged on a widefield microscope. Stills from a movie where metaphase cells were each treated with rapamycin 200 nM are shown. The post-rapamycin image is ~15 min after treatment. Scale bar, 10 μm.

### Defining mitotic clathrin, TACC3, chTOG and GTSE1 interactions using knocksideways of endogenously tagged proteins

Acute manipulation of a protein localization using knocksideways can be used to uncover interactions in living cells (Hood et al., 2013). To examine mitotic interactions between clathrin, TACC3, chTOG and GTSE1, we set out to relocalize each endogenous protein in mitotic knock-in cells and ask whether this manipulation affects the localization of the other proteins, detected by indirect immunofluorescence (Figure 3). Relocalization of endogenous clathrin (CLTA-FKBP-GFP) caused the removal of TACC3, GTSE1 but not chTOG from the spindle (Figure 3A). Similarly, relocalization of endogenous GFP-FKBP-TACC3 resulted in removal of clathrin (very small effect) and GTSE1, but not chTOG from the spindle (Figure 3B). In contrast, relocalization of either chTOG-FKBP-GFP or GTSE1-FKBP-GFP had no effect on the spindle localization of the other three respective proteins (Figure 3C,D). While these experiments were designed to examine interactions between endogenous proteins, the use of immunofluorescence had two drawbacks. Firstly, antibody specificity was an issue. No effect on chTOG localization was seen after relocalization of each of the four proteins, including chTOG-FKBP-GFP itself; and relocalization of GTSE1-FKBP-GFP was not detected by anti-GTSE1. Secondly, single cells could not be tracked during knocksideways.

**Figure 3.**
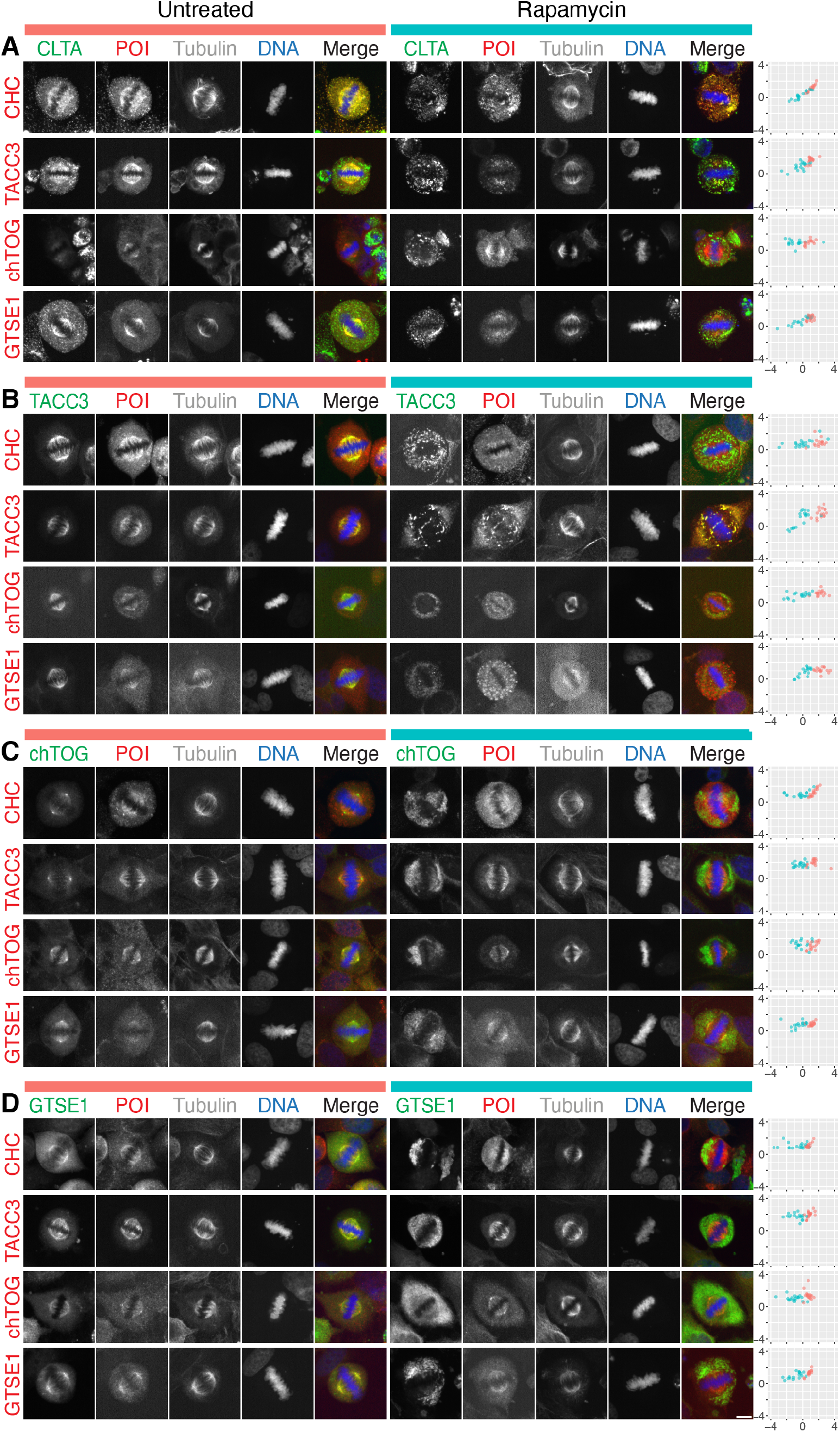
Co-rerouting of endogenous complex members following knocksideways in knock-in cell lines. Knocksideways experiments using each knock-in cell line expressing dark MitoTrap. (**A**) CLTA-FKBP-GFP cells. (**B**) GFP-FKBP-TACC3 cells. (**C**) chTOG-FKBP-GFP cells (**D**) GTSE1-FKBP-GFP cells. Representative widefield micrographs of cells that were treated with rapamycin (200 nM) for 30 min,fixed and stained for tubulin and either CHC, TACC3, chTOG, or GTSE1 (protein-of-interest, POI, red). Scale bar, 10 μm. Right, quantification of images. Spindle localization of the target protein (x-axis) and the protein-of-interest (y-axis) in control (salmon) and knocksideways (turquoise) cells. Spindle localization is the ratio of spindle to cytoplasmic fluorescence shown on a log2 scale (1 is twice the amount of protein on the spindle as the cytoplasm, −1 indicates half the amount on spindle versus cytoplasm). Quantification of cells from three or more experiments are shown.

We next sought to repeat these experiments using a live cell approach. To do this, the knock-in cell lines were transfected with dark mitotrap and either mCherry-CLTA, mCherry-TACC3, chTOG-mCherry or tdTomato-GTSE1. Metaphase cells were imaged live as rapamycin was added (Figure 4). Relocalization of endogenous clathrin caused the removal of mCherry-TACC3, chTOG-mCherry and tdTomato-GTSE1 from the spindle (Figure 4A). Similarly, relocalization of endogenous TACC3 also caused the removal of the other three proteins from the spindle but to a lesser extent than with clathrin (Figure 4B). Again, relocalization of either chTOG or GTSE1 had no effect on the spindle localization of the other three respective proteins (Figure 4C,D). A semi-automated analysis procedure was used to measure induced relocalization of both proteins (see Methods). All movement was from the mitotic spindle to the mitochondria, without significant l oss t o t he cytoplasm, suggesting that the complex is either relocalized *en masse* or not. 2D arrow plots were therefore used to visualize the results of these experiments (Figure 4). As previously reported, mCherry-TACC3 expression distorted the localization of the complex prior to knocksideways (Booth et al., 2011; Nixon et al., 2015), enhancing the amount of clathrin, chTOG and GTSE1 on the spindle (Figure 4, note the rightward shift of the starting point in the arrow plots when mCherry-TACC3 was expressed). This likely reflects the importance of TACC3 in loading the complex onto the spindle (Hood et al., 2013).

**Figure 4.**
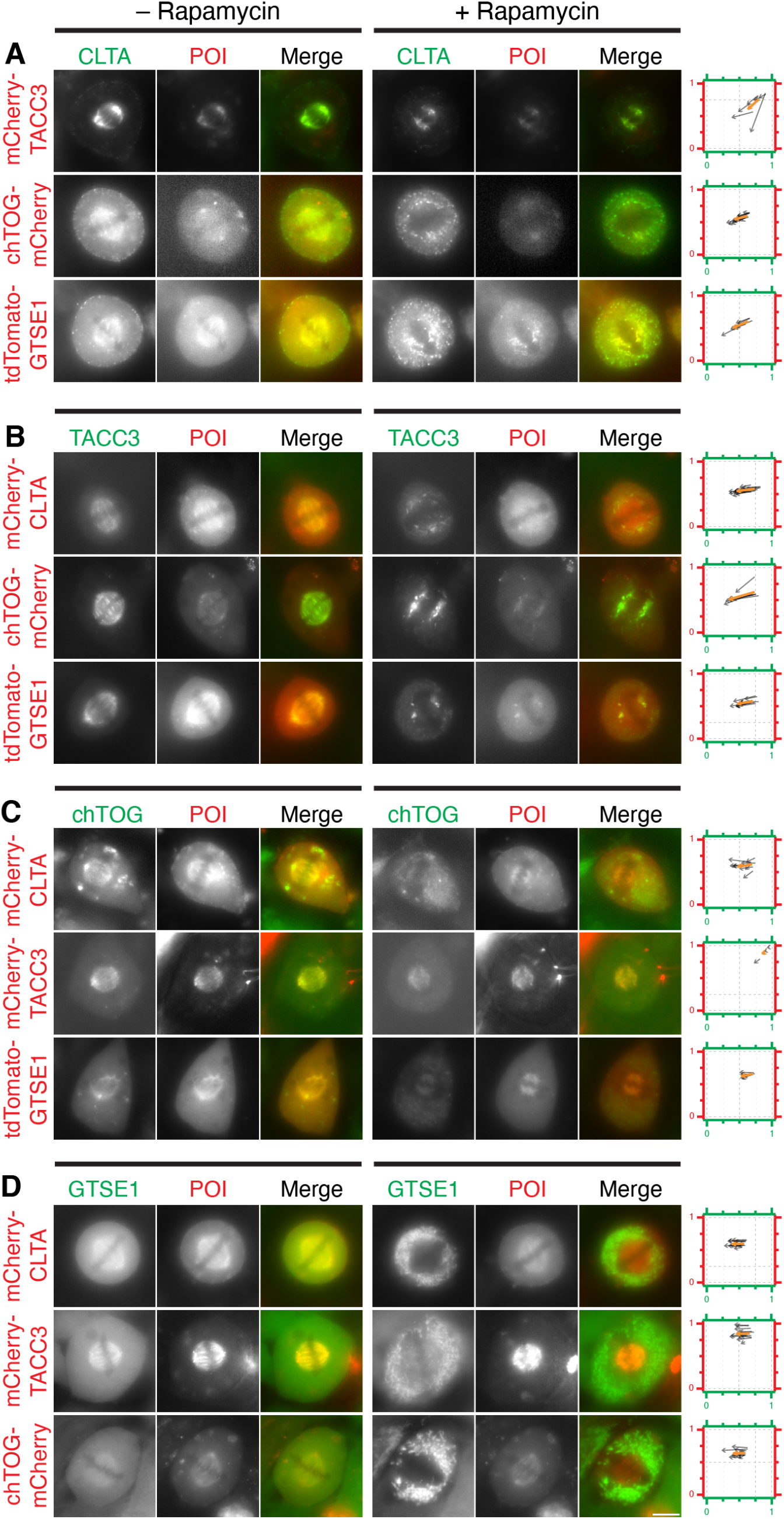
Co-rerouting of complex members during live-cell knocksideways experiments in knock-in cell lines. Knocksideways experiments using each knock-in cell line expressing dark MitoTrap and one of the three other complex proteins tagged with a red fluorescent protein (mCherry-CLTA, mCherry-TACC3, chTOG-mCherry or tdTomato-GTSE1) as indicated. (**A**) CLTA-FKBP-GFP cells. (**B**) GFP-FKBP-TACC3 cells. (**C**) chTOG-FKBP-GFP cells. (**D**) GTSE1-FKBP-GFP cells. Still images are shown before and 10 min after rapamycin (200 nM) treatment. Scale bar, 10 μm. Right, arrow plots show extent of co-rerouting. Arrows show the fraction of spindle and mitochondria fluorescence that is at the spindle (i.e. 1 = completely spindle-localized, 0 = mitochondria-localized), for both channels, moving from pre to post rapamycin localization. Black arrows represent individual cells measured across three experimental repeats, orange arrow indicates the mean.

The expression of other partner proteins mCherry-CLTA, chTOG-mCherry, and tdTomato-GTSE1 had no effect on the localization of the knock-in protein.

The results of both knocksideways approaches are summarized in Supplementary Table 2. Relocalization of either clathrin or TACC3 during metaphase results in removal of the entire TACC3–chTOG–clathrin–GTSE1 complex. The efficiency of this removal is higher with clathrin than TACC3. Relocalization of either chTOG or GTSE1 has no effect on the rest of the complex, suggesting that these proteins are ancillary to TACC3–chTOG–clathrin–GTSE1.

### Role of LIDL motifs in recruitment of GTSE1 to the TACC3–chTOG–clathrin complex

In order to test if GTSE1 is an ancillary complex member, we sought to disrupt its interaction with clathrin and assess whether or not the spindle-binding of these two proteins was interdependent. To examine the effect on the mitotic localization of both proteins, mCherry-tagged GTSE1 constructs were expressed in GTSE1-depleted CLTA-FKBP-GFP cells (Figure 5). GTSE1 has a previously-mapped clathrin interaction domain (CID, 639-720), containing five c lathrin b ox-like m otifs LI[DQ][LF] (hereafter referred to as LIDL motifs), which was targeted for disruption (Wood et al., 2017; Rondelet et al., 2020). We found that deletion of the entire CID resulted in a reduction in GTSE1 on the spindle. Mutation of LIDL motifs 1-2, 3 or 4-5 to alanines did not result in reduction, but when mutated in combination resulted in a loss of GTSE1 that was similar to deletion of the CID. However, under all conditions the spindle localization of clathrin was unaffected.

**Figure 5.**
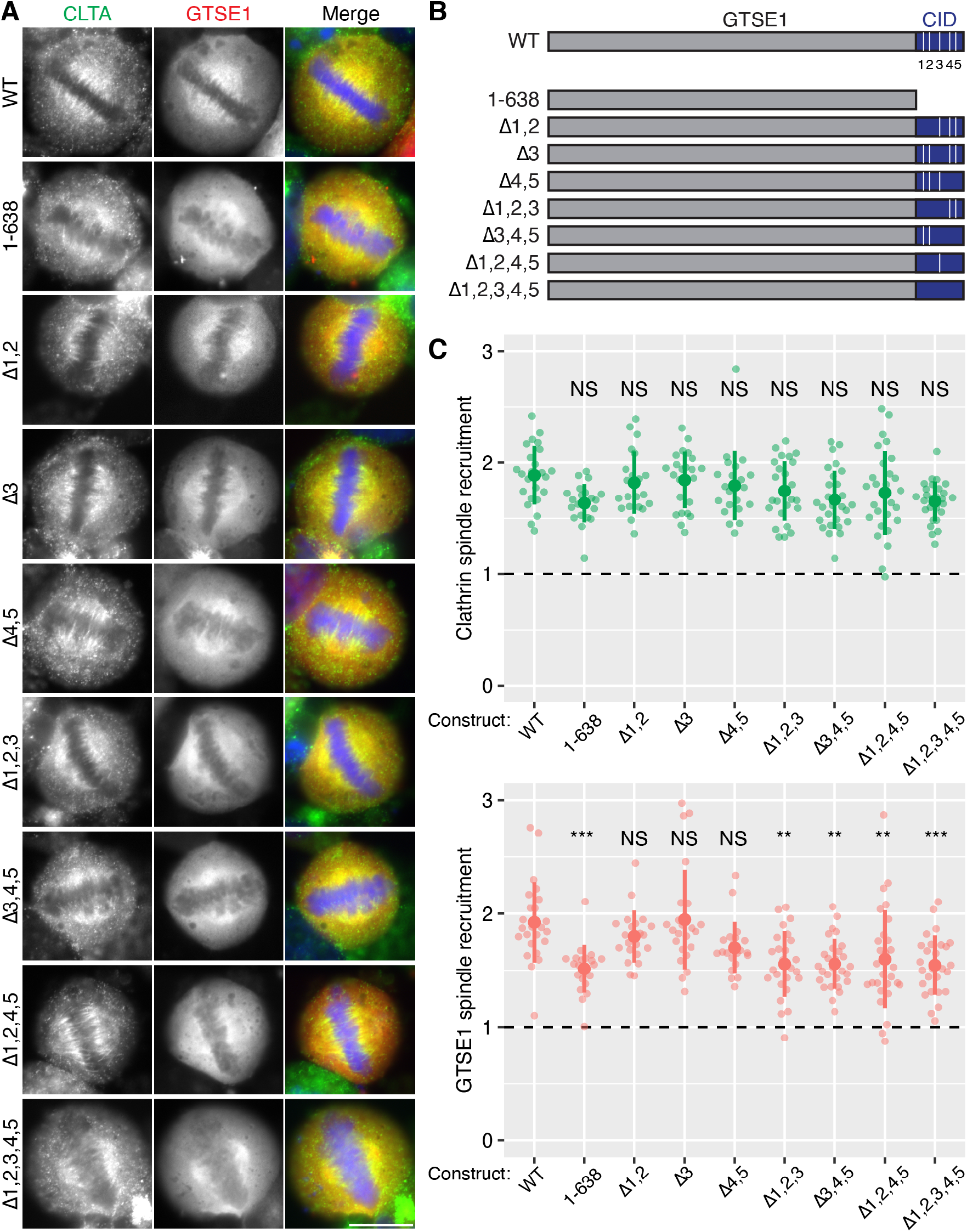
Role of LIDL motifs in GTSE1 spindle localization. (**A**) Representative widefield micrographs of GTSE1-mCherry constructs (red) in GTSE1-depleted CLTA-FKBP-GFP cells at metaphase. Cells expressing the indicated construct were fixed and stained using DAPI (blue) and a GFP-boost antibody to enhance the signal of CLTA-FKBP-GFP (green). Scale bar, 10 μm. (**B**) Schematic diagram of full-length GTSE1 (WT, 1-720) and mutant forms. The five LIDL motifs are denoted 1 through 5. Mutation of the corresponding motif by the introduction of four alanine residues is denoted by Δ. (**C**) Quantification of the spindle localization of clathrin and GTSE1. Each dot represents a single cell, n = 21-28 cells per construct over 3 separate experiments. The dashed horizontal line represents no enrichment on the spindle. The large dot and error bars show the mean ±SD, respectively. Analysis of variance (ANOVA) with Tukey’s post-hoc test was used to compare the means between each group. The p-value level is shown compared to WT: ***, *p* < 0.001; **, *p* < 0.01; NS, *p* > 0.05.

To test if the reduction in GTSE1 spindle localization represented a block of recruitment, cells were treated with 0.3 μM MLN8237 to inhibit Aurora A activity and provide a reference for minimal recruitment (Hood et al., 2013; Booth et al., 2011). Spindle localization of both clathrin and GTSE1 WT was abolished by drug treatment (Figure S3). Again, spindle localization of GTSE1Δ1,2,3,4,5 was lower than WT in untreated cells, and was not reduced further by MLN8237 treatment (*p* = 0.08). These data are consistent with the idea that GTSE1 is recruited to the spindle by clathrin via multiple LIDL motifs in GTSE1 (Rondelet et al., 2020). Moreover they suggest that there is no interdependent spindle localization of clathrin-GTSE1 and that GTSE1 is an ancillary member of the complex.

The ability to bind clathrin is necessary for GTSE1 to localize to the spindle, but is it sufficient? To address this question we examined the subcellular localization of a panel of GTSE1 fragments in mitosis or interphase cells (Figure 6). A GTSE1 fragment comprising the CID containing all five LIDL motifs was unable to bind the mitotic spindle. Progressively adding more N-terminal sequence to the CID eventually yielded a construct that bound the spindle (161-720, Figure 6A-C). This experiment demonstrated that the CID alone is not sufficient for spindle localization. Interphase microtubule-binding was seen for the GTSE1 fragment 161-720 and to a lesser extent for 1-354, 335-720 and 400-720 (Figure 6A,D). This suggests that the region 161-638 contains one or more regions that can bind microtubules and that these regions, together with the 5XLIDL motifs in the CID are required for spindle localization.

**Figure 6.**
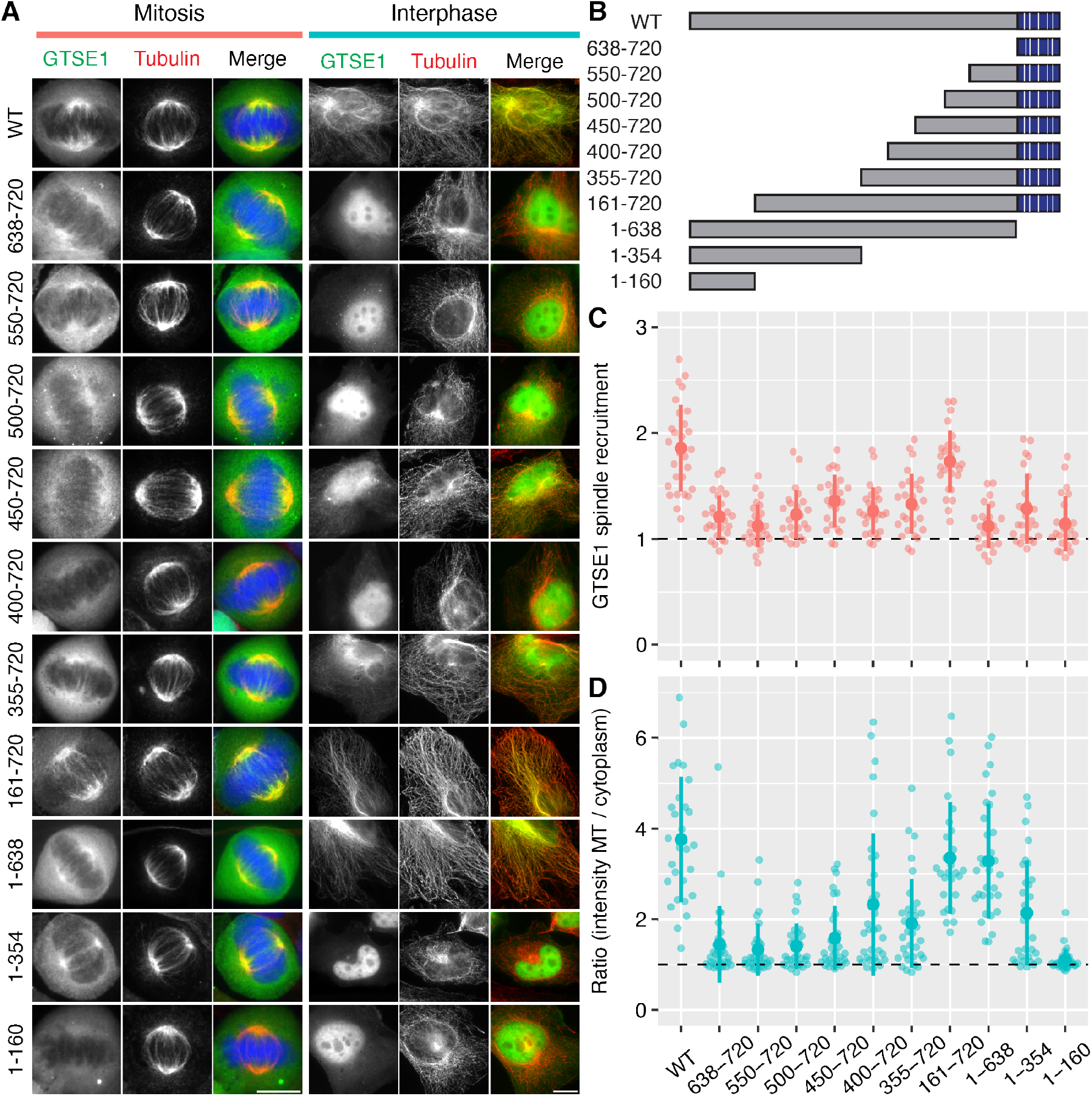
Localization of GTSE1 fragments in interphase and mitosis. (**A**) Representative widefield micrographs of GTSE1-FKBP-GFP constructs expressed in cells in mitosis or interphase. Scale bar, 10 μm. (**B**) Schematic diagram of full-length GTSE1 (WT, 1-720) and fragments of GTSE1 used in this figure. Quantification of GTSE1 localization on mitotic spindles (**C**) or interphase microtubules (**D**). Each dot represents a single cell, n = 23-28 cells per construct (mitosis) and n = 27-33 cells per construct (interphase) pooled from three independent experiments. The large dot and error bars show the mean and the mean ±SD, respectively. Analysis of variance (ANOVA) with Tukey’s post-hoc test was used to compare the means between each group. The p-value level is shown compared to WT: ***, *p* < 0.001; NS, *p* > 0.05.

### PIK3C2A is not a component of the TACC3–chTOG–clathrin–GTSE1 complex

We next investigated whether or not PIK3C2A is a component of the TACC3–chTOG–clathrin–GTSE1 complex, since PIK3C2A has been proposed to bind TACC3 and clathrin, and therefore stabilize the complex (Gulluni et al., 2017). If PIK3C2A binds the complex, we would predict that it should also localize to the mitotic spindle. We imaged GFP-PIK3C2A in live cells and found no evidence for spindle localization (Figure 7A). The construct localized to clathrin-coated pits suggesting that the GFP-tag had not interfered with its normal localization. We next overexpressed mCherry-TACC3 to concentrate the TACC3–chTOG–clathrin–GTSE1 complex on the spindle and maximize our chances of seeing any GFP-PIK3C2A signal on microtubules, but again, we saw no spindle localization of GFP-PIK3C2A (Figure 7B).

**Figure 7.**
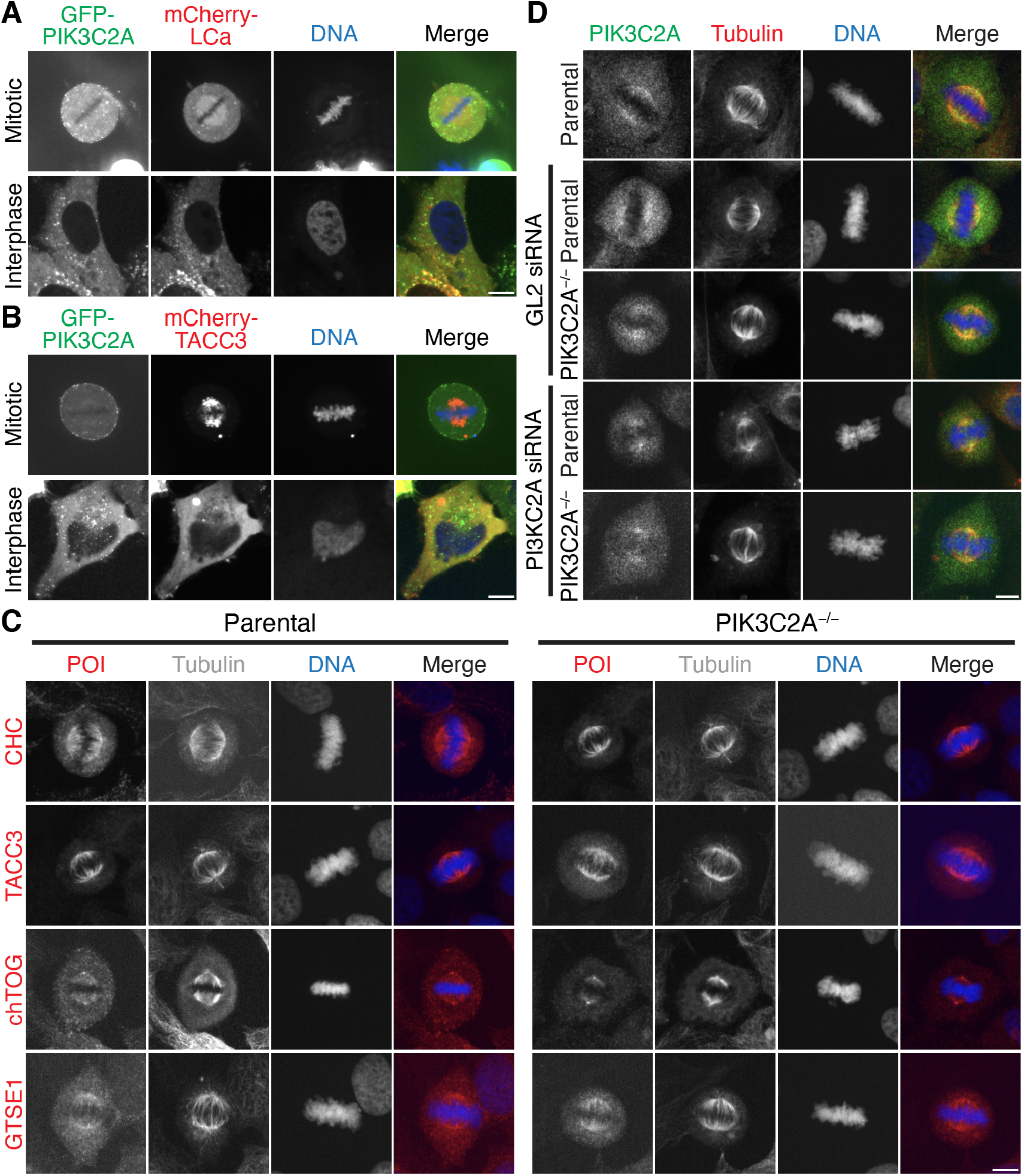
PIK3C2A is not a component of the TACC3–chTOG–clathrin–GTSE1 complex. Representative confocal micrographs of HeLa cells expressing GFP-PIK3C2A and either mCherry-CLTA (**A**) or mCherry-TACC3 (**B**). (**C**) Representative widefield micrographs of parental HeLa and PIK3C2A^−/−^ cells stained for tubulin and either CHC, TACC3, chTOG, or GTSE1 (red). (**D**) Representative widefield micrographs of parental HeLa and PIK3C2A^−/−^ cells treated with GL2 (control) or PIK3C2A siRNA, stained with an anti-PIK3C2A antibody (Proteintech, Green) and tubulin (red). Scale bar, 10 μm.

To further explore any mitotic role for PIK3C2A, we generated a PIK3C2A knockout cell line using CRISPR/Cas9. This generated a clone with a premature stop codon in both alleles, resulting in 1-87 and 1-72 residues, that we termed PIK3C2A null (Supplementary Figure S4C). It was previously shown that PIK3C2A knockout in primary mouse embryo fibroblasts (MEFS) altered their mitotic progression (Gulluni et al., 2017). We analyzed mitotic timings of our PIK3C2A null cell line, compared to parental HeLa cells, and found no differences in mitotic timings (Supplementary Figure S4D).

If PIK3C2A was a scaffold protein for the TACC3–chTOG–clathrin–GTSE1 complex, we would expect some disruption of the spindle localization of clathrin, TACC3, chTOG or GTSE1 in the PIK3C2 null cells. However, immunostaining of parental HeLa and PIK3C2A null cells with antibodies against CHC, TACC3, chTOG and GTSE1, revealed a similar distribution of all four complex members during mitosis (Figure 7C).

In the original paper, immunostaining of PIK3C2A at the mitotic spindle was shown (Gulluni et al., 2017). We immunostained parental HeLa and the PIK3C2A-null cells with the same anti-PI3KC3A antibody used in the original report and found that there was a signal at the mitotic spindle, but that it was non-specific since it was also detected in the PIK3C2A-null cells (Figure 7D). We also used RNAi of PIK3C2A in parental and PIK3C2A-null cells to rule out the possibility that the antibody signal resulted from residual expression of PIK3C2A. Again, the spindle fluorescence remained after RNAi indicating that the antibody is non-specific for immunofluorescence. Taken together, our results suggest that PIK3C2A is not a component of the TACC3–chTOG–clathrin–GTSE1 complex.

## Discussion

Inducible relocalization is a powerful method to investigate protein-protein interactions in cells and to pinpoint where and when they occur. We generated a number of cell lines to study the interactions between members of the TACC3–chTOG–clathrin–GTSE1 complex on mitotic spindles at metaphase. This approach showed that TACC3 and clathrin are core complex members, while chTOG and GTSE1 are ancillary. Our current picture of this multiprotein complex is outlined in Figure 8.

**Figure 8.**
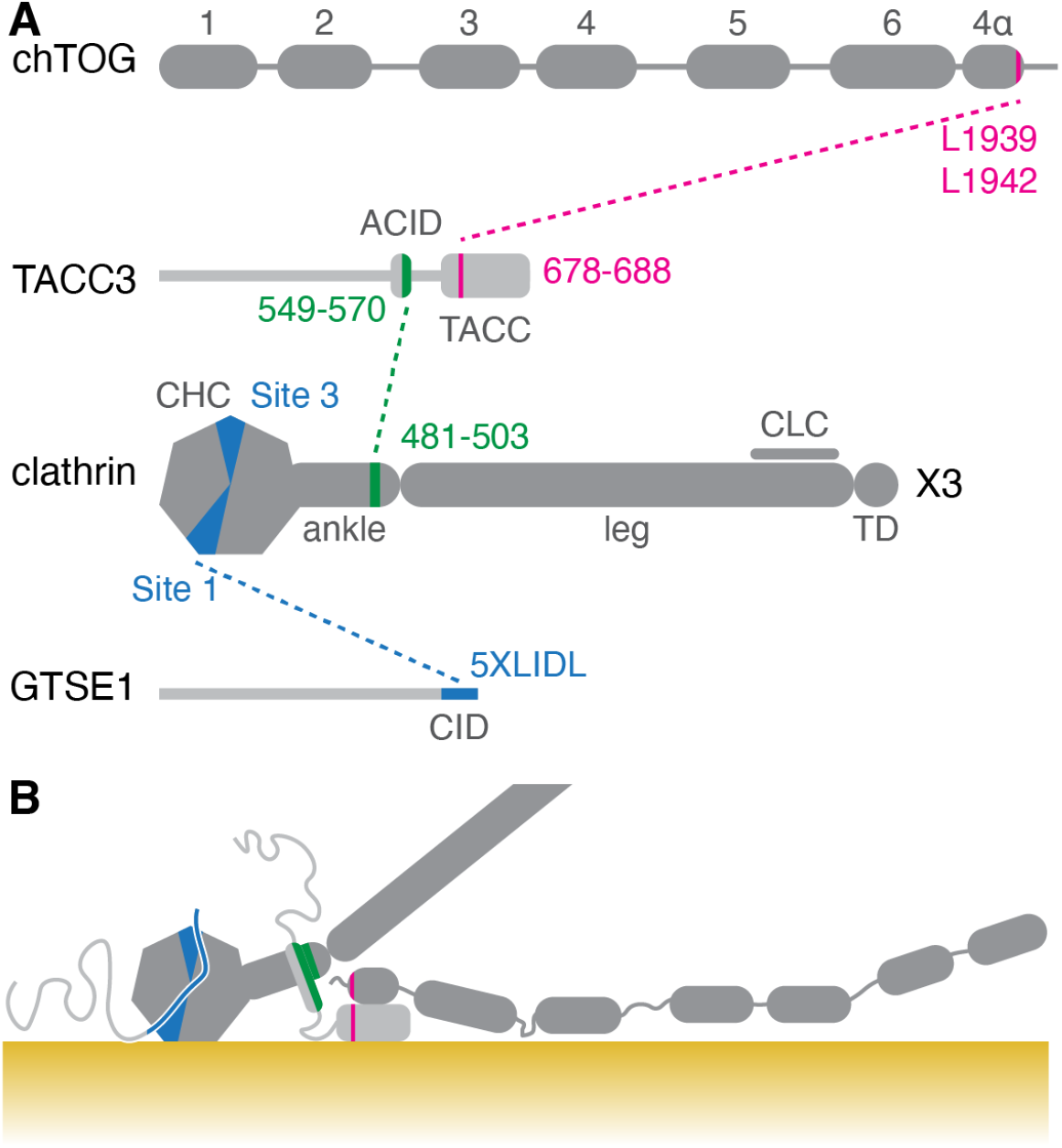
Summary diagram of interactions between TACC3–chTOG–clathrin–GTSE1 complex members and microtubules. (**A**) Primary structure of chTOG, TACC3, clathrin and GTSE1, showing the interactions between each protein. Abbreviations: ACID, Aurora-A and clathrin interaction domain; CHC, clathrin heavy chain; CLC, clathrin light chain; TD, trimerization domain. Interactions were mapped previously (Gutiérrez-Caballero et al., 2015; Hood et al., 2013; Burgess et al., 2018; Rondelet et al., 2020). (**B**) Proposed topology of the complex on a microtubule (yellow). TACC3 and clathrin bind each other and form a composite microtubule interaction surface. GTSE1 and chTOG bind to clathrin and TACC3, respectively. Both proteins can interact with microtubules: chTOG in a domain between TOG4 and TOG5 (Widlund et al., 2011), GTSE1 in a diffuse region between residues 161-638 (this work) although neither interaction is necessary for the complex to bind microtubules.

It had been reported that PIK3C2A is a component of the TACC3–chTOG–clathrin–GTSE1 complex, where it was proposed to act as a scaffold protein binding both TACC3 and clathrin (Gulluni et al., 2017). This proposal was consistent with several observations. First, PIK3C2A was found to interact with clathrin, GTSE1 and TACC3 in a proteomic analysis of immunoprecipitations with each of these three proteins from mitotic lysate (Hubner et al., 2010). In that study, immunoprecipitation of PIK3C2A brought down clathrin, GTSE1 and components of the membrane trafficking machinery, but notably neither TACC3 nor chTOG co-immunoprecipitated with PIK3C2A. Second, PIK3C2A binds clathrin heavy chain via an N-terminal region which contains a clathrin box-like motif (LLLDD) (Gaidarov et al., 2001). These motifs bind to grooves in the seven-bladed beta-propeller that constitutes the N-terminal domain of clathrin heavy chain (Smith et al., 2017). The N-terminal domain is required for clathrin–TACC3 to localize at the spindle (Royle et al., 2005) and mutations in one of the grooves is sufficient to reduces pindle-binding (Hood et al., 2013). However, while the proposal that PIK3C2A was a component of the complex made sense, we found no evidence to suggest that PIK3C2A was even present on mitotic spindles. GFP-tagged PIK3C2A was found in clathrin-coated vesicles as expected, but was absent from the mitotic spindles of HeLa cells. We also found that the PIK3C2A antibody used in the original study to detect the protein at the spindle gave a false signal that remained after knockout and/or knockdown of PIK3C2A. Finally, a PIK3C2A-null cell line we generated had no mitotic delays, and all members of the TACC3–chTOG–clathrin–GTSE1 complex had normal localization. We conclude that PIK3C2A is not a component of the complex and any mitotic function for this protein is doubtful. PIK3C2A has a well-established role in clathrin-mediated membrane traffic (Posor et al., 2013) and it seems likely that the presence of PIK3C2A among other membrane trafficking factors in the original proteomic work was due to association with a fraction of clathrin that was not associated with the spindle, or erroneous binding during purification (Hubner et al., 2010).

Recent work has shown that GTSE1 contains five conserved LIDL motifs an intrinsically disordered C-terminal region which can bind to the N-terminal domain of clathrin heavy chain (Rondelet et al., 2020). In agreement with this, we found that these motifs are redundant and that mutations which reduce the total number of motifs to below three, impaired spindle-binding significantly. We also found that GTSE1 was an ancillary protein not required for the localization of the complex on microtubules and that inducing its mislocalization did not affect the other complex members, this interpretation is consistent with other work on GTSE1 (Rondelet et al., 2020; Bendre et al., 2016). It is a mystery why mutation of one groove of the N-terminal domain of clathrin heavy chain results in loss of the complex from the spindle, since it appears that this domain recruits GTSE1 to k-fibers but that GTSE1 is not needed for localization (Hood et al., 2013). One explanation is that this groove interacts directly with microtubules and GTSE1 may also bind other sites on clathrin’s N-terminal domain. Another is that the GTSE1–clathrin interaction may be important for the formation of the complex, but not for its stability once loaded onto microtubules.

The ancillary nature of GTSE1 and chTOG binding to the complex via association with clathrin and TACC3 respectively is intriguing. Especially since GTSE1 and chTOG each have the ability to bind microtubules themselves (Monte et al., 2000; Spittle et al., 2000). Rondelet et al. proposed that clathrin–TACC3 could be forming a “scaffold” for the recruitment of other factors, such as GTSE1, to the spindle so that they can in turn perform specific functions (Rondelet et al., 2020; Bendre et al., 2016). Our work is consistent with this idea, that clathrin–TACC3 are core to spindle microtubule-binding and that other ancillary factors may be recruited through this complex. In this work we mapped a constitutive microtubule-binding region in GTSE1 to residues 161-638, while chTOG likely binds the microtubule lattice through a region between TOG4 and TOG5. The criterion for binding clathrin–TACC3 at the spindle may include the ability to bind microtubules, which would explain the selectivity for ancillary partners and mean that clathrin adaptors for example are not recruited to the spindle.

Our work establishes that, in order to disrupt the TACC3–chTOG–clathrin–GTSE1 complex, agents that target i) the TACC3–clathrin interaction or ii) the interface between TACC3–clathrin and microtubules are required. In the first case, preventing the helix that is formed by phosphorylation of TACC3 on serine 558 from binding to the helical repeat in the ankle region of clathrin heavy chain is predicted to disrupt the complex (Burgess et al., 2018). Second, the microtubule interface needs to be mapped at high resolution using cryo-electron microscopy. The endogenously tagged cell lines we have developed will be useful for pursuing these questions. Besides fluorescence microscopy and knocksideways, the cells are well suited for visualizing proteins at the ultrastructural level using inducible methodologies such as FerriTagging (Clarke and Royle, 2018).

## Methods

### Molecular biology

The following plasmids were available from previous work: Mitotrap (pMito-mCherry-FRB), dark Mitotrap (pMito-mCherryK70N-FRB), mCherry-alpha-Tubulin, mCherry-CLTA (mCherry-LCa), mCherry-TACC3, and chTOG-GFP (Cheeseman et al., 2013; Wood et al., 2017; Hood et al., 2013). Plasmid to express chTOG-mCherry was made by ligating a BamHI-NotI fragment from chTOG-GFP into pmCherry-N1. For tdTomato-GTSE1, a GFP-GTSE1 construct was first made by PCR of full-length human GTSE1 (IMAGE: 4138532) with the addition of EcoRI-BamHI and cloning into pEGFP-C1 and then ligating EcoRI-BamHI fragment from GFP-GTSE1 into ptdTomato-C1. For GFP-PIK3C2A a ScaI-BstEII fragment from full-length human PIK3C2A in PCR-XL-Topo (IMAGE:8322710) was cloned into pEGFP-C1. The mCherry-tagged GTSE1 constructs were made by PCR amplification o f G TSE1 (IMAGE: 4138532) followed by insertion into pmCherry-N1 between SalI-BamHI, and using site-directed mutagenesis to introduce each mutation.

### Cell culture

HeLa cells (Health Protection Agency/European Collection of Authenticated Cell Cultures, #93021013) were maintained in Dulbecco’s Modified Eagle’s Medium (DMEM) supplemented with 10 % FBS and 100 Uml^−1^ penicillin/streptomycin in a humidified incubator at 37 °C and 5 % CO_2_.

Knock-in cell lines were generated by CRISPR/Cas gene editing. The orientation of tags (N-terminal or C-terminal) was guided by previous work on CLTA (Doyon et al., 2011), TACC3 (Cheeseman et al., 2013), chTOG (Gutiérrez-Caballero et al., 2015), and GTSE1 (Scolz et al., 2012). Briefly, HeLa cells were transfected with a Cas9n D10A nickase plasmid (pSpCas9n(BB)-2A-Puro, pX462, Addgene #48141) and a repair template. The following guide pairs were used; CLTA-FKBP-GFP cell line (guide 1, 5’-CACCGCAGATGTAGTGTTTCCACA-3’; guide 2, 5’-CACCGTGAAGCTCTTCACAGTCAT-3’), GFP-FKBP-TACC3 cell line (guide 1, 5’-CACCGGCACGACCACTTCCCACAC-3’; guide 2, 5’-CACCGACGTCTGTGTCTGGACAATG-3’), chTOG-FKBP-GFP cell line (guide 1, 5’-CACCGAAGATCCTCCGACAGCGATG-3’; guide 2, 5’-CACCGCCAGACCACATCGCTGTCGG-3’), FKBP-GFP-GTSE1 cell line (guide 1, 5’-CACCGGGAGCTCAGGTCTATGAGC-3’; guide 2, 5’-CACCGTGAGGCTGACAAGGAGAACG-3’). Details of the repair templates are available (see below). Ten days after transfection, single GFP-positive cells were selected by fluorescence-activated cell sorting (FACS), expanded and validated by microscopy, Western blotting, PCR and DNA sequencing.

The PIK3C2A knockout cell line was generated by transfecting HeLa cells with pSpCas9(BB)-2A-GFP (pX458, Addgene #48138) into which a single guide (5’-CACCGAGCACAGGTTTATAACAAGC-3’) had been cloned. GFP-positive cells were isolated by FACS and then single cell clones were validated by western blotting and genome sequencing. Briefly, a genomic region encompassing the target site was amplified (forward, 5’-CCAGTTGTGTCAGGAAATGGG-3’; reverse, 5’-TCCAAATCAGTCCTTGCTTTCCC-3’) and TA-cloned into pGEM-T Easy vector. Ten bacterial transformants were picked and sequenced, revealing a 1:1 ratio of the two alleles shown in Supplementary Figure S4.

For knockdown of endogenous GTSE1 in spindle recruitment experiments, CLTA-FKBP-GFP CRISPR knock-in HeLa cells were transfected with 100 nM siRNA targeting the 3’UTR of GTSE1 (GTSE1, 5’-GCCTGGGAAATATAGTGAAACTCCT-3’; GL2 (control), 5’-CGTACGCGGAATACTTCGA-3’). For knockdown of PIK3C2A in HeLa RNAi was performed by transfecting 60 nM siRNA (siCtrl ‘GL2’ 5’-CGTACGCGGAATACTTCGA-3’; siPIK3C2A ‘1’ 5’-GAAACTATTGCTGGATGACAGT-3’), using Lipofectamine 2000 (Invitrogen), according to the manufacturer’s instructions.

For DNA plasmid transfections, cells were transfected with a total of 1000 μg to 1500 μg DNA in 35 mm fluorodishes or 6 well plates using Genejuice as per the manufacturer’s instructions (Merck Millipore). Cells were typically imaged 48 h after transfection. For knocksideways experiments, cells were transfected with plasmids to express MitoTrap alone or Dark MitoTrap in combination other constructs as indicated. For the expression of GTSE1-mCherry mutants, cells were transfected with DNA 24 h after siRNA treatment using Genejuice (Merck Millipore), following the manufacturer’s protocol. Cells were fixed 64 h after siRNA transfection and 40 h after DNA transfection.

Knocksideways was via the application of rapamycin (Alfa Aesar) to a final concentration of 200 nM; either live on the microscope or, in the case of immunofluorescence experiments, for 30 min prior to fixation. For Aurora-A inhibition, MLN8237 (Apexbio) was added at a final concentration of 300 nM for 40 min.

### Immunofluorescence

For immunofluorescence, cells were fixed at RT using PTEMF (20 mM Pipes, pH 6.8, 10 mM EGTA, 1 mM MgCl_2_, 0.2 % Triton X-100, and 4 % paraformaldehyde) for 10 min and permeabilized at RT in 0.5 % Triton-X100 in PBS for 10 min. Cells were blocked in 3 % BSA in PBS for 30 min. Cells were incubated for 1 h at RT with primary antibodies as follows: rabbit anti-Tubulin (PA5-19489, Invitrogen), mouse anti-Tubulin (B-5-1-2, Sigma), mouse anti-CHC (X22, CRL-2228 ATCC), rabbit anti-chTOG (34032, QED Biosciences), mouse anti-TACC3 (ab56595, Abcam), mouse anti-GTSE1 (H00051512-B01P, Abnova), rabbit anti-PIK3C2A (22028-1-AP, Proteintech). Cells were washed three times with PBS, then incubated for 1 h with Alexa568- or Alexa647-conjugated secondary antibodies (Molecular Probes). Finally, cover slips were rinsed with PBS and mounted with mowiol containing DAPI. In some experiments it was necessary to boost the GFP signal of the knock-in cells. To do this, GFP-booster (Alexa488, Chromotek) or GFP Rabbit anti-Tag, Alexa Fluor 488 (Invitrogen) 1:200 was used during the primary incubations.

### Biochemistry

For Western blotting, cell lysates were prepared by scraping cells in RIPA buffer containing cOmplete EDTA-free protease inhibitor cocktail (Sigma-Aldrich), incubation on ice for 30 min, and clarification in a benchtop centrifuge for 15 min at 4 °C. Lysates were boiled in 4× Laemmli buffer for 10 minute and resolved on a precast 4 to 15 % polyacrylamide gel (Bio-Rad). Proteins were transferred to nitrocellulose using a Trans-Blot Turbo Transfer System (Bio-Rad). Primary antibodies used were mouse anti-alpha tubulin (DM1A, Sigma) 1:10,000, rabbit anti-CLTA (sc-28276, Santa Cruz) 1:1000, goat anti-TACC3 (AF5720, Novus Biologicals) 1 μg ml^−1^, rabbit anti-chTOG (34032, QED Biosciences) 1:2000, mouse anti-GTSE1 (H00051512-B01P, Abnova) 1:500 and rabbit anti-PIK3C2A (22028-1-AP, Proteintech) 1:1000. Secondary antibodies used were mouse, rabbit and goat HRP conjugates. For detection, enhanced chemiluminescence detection reagent (GE Healthcare) and manual exposure of Hyperfilm (GE Healthcare) was performed.

### Microscopy

For live cell imaging, media was changed to Leibovitz (Gibco) L-15 CO_2_-independent medium supplemented with 10 % FBS. Imaging was performed on a Nikon Ti epifluorescence microscope with standard filter sets and 100× or 60× (both 1.4 NA, oil, PlanApoVC) objectives, equipped with a heated environmental chamber (OKOlab) and CoolSnap MYO camera (Photometrics) using NIS elements AR software.

For overnight mitotic imaging, cells were incubated with 0.5 μM Sir-DNA (Spirochrome) for 60 min to visualize DNA. Image stacks (7 × 2 μm optical sections; 1 × 1 binning) were acquired every 3 min for 12 h with a 40× oil-immersion 1.3 NA objective using an Olympus DeltaVision Elite microscope (Applied Precision, LLC) equipped with a CoolSNAP HQ2 camera (Roper Scientific). Images were acquired at 10 % neutral density using Cy5 filter and an exposure time of 100 ms. A stage-top incubator maintained cells at 37 °C and 5 % CO_2_ with further stabilization from a microscope enclosure (Weather station, PrecisionControl) held at 37 °C. To image fixed cells, image stacks (5 × 1 μm optical sections) were acquired on a spinning disc confocal system (Ultraview Vox, Perkin Elmer) with a 60× 1.4 NA oil-immersion objective (Nikon) and Hamamatsu Orca-R2 camera.

### Data Analysis

Analysis of knocksideways movies was done by extracting a pre- and a post-rapamycin multi-channel image from the sequence. An automated procedure in Fiji measured three regions in each of the following areas: spindle, cytoplasm and mitochondria, after registration of the pre and post images. A background measurement and a whole cell fluorescence measurement were also taken. The average value for each region, after background subtraction, was corrected for bleach using the whole cell fluorescence measurement (background-subtracted) for the respective channel. Data were exported as csv and read into IgorPro (WaveMetrics) where a custom-written procedure analyzed the data and generated all the plots. Ternary diagrams of spindle, mitochondria and cytoplasm fluorescence revealed that knocksideways resulted in movement mainly between spindle and mitochondria (Supplementary Figure S5)). Therefore, the fraction of fluorescence at the spindle and mitochondria were used to generate the arrow plots.

For spindle localization analysis of fixed cells, a 31 × 31 pixel (1.4 μm^2^) ROI was used to measure three regions of the spindle, the cytoplasm and one region outside of the cell as background, using Fiji. Following background-subtraction, the average spindle fluorescence was divided by the cytoplasm fluorescence to give a measure of spindle enrichment. To quantify the MT localization of GTSE1 fragments, a line-scan analysis method adapted from (Hooikaas et al., 2019) was used. Using an automated procedure in Fiji, average fluorescence intensities from three lines, 1 μm to 3 μm length, along MTs stained for *α*-tubulin and three adjacent lines (not coincident with MTs) were measured. Following background subtraction, the average fluorescence intensity of the MT line scan was divided by the average fluorescence intensity of the adjacent control line scan to generate a MT enrichment ratio. Analysis was done by an experimenter blind to the conditions of the experiment.

All figures were made in Fiji, R or Igor Pro 8 and assembled using Adobe Illustrator.

### Data and software availability

All code used in the manuscript and sequences for repair templates is available at https://github.com/quantixed/p053p030.

## ACKNOWLEDGEMENTS

We thank Computing and Advanced Microscopy Unit (CAMDU) for their support and assistance in this work. We are also grateful to Nuria Ferrándiz, Vishakha Karnawat and Laura Downie for commenting on the manuscript. The work was supported by a Programme Award from Cancer Research UK (C25425/A27718). JS was supported by a studentship from BBSRC Midlands Integrative Biosciences Training Partnership (MIBTP) (BB/M01116X/1).

The authors declare no conflict of interest.

## Author contributions

E.L. Ryan did the majority of the experimental work and wrote the first draft of the manuscript. J. Shelford contributed the GTSE1 experiments, generated the GTSE1 cell line, helped to validate TACC3 knock-in cells, and wrote R scripts for data analysis. T. Massam-Wu helped to generate and validate knock-in cell lines. R. Bayliss helped supervise the project. S.J. Royle supervised the work, wrote computer code, and contributed to figure and paper preparation.

## Supplementary Information

**Table 1.** Summary of knock-in cell lines used in this study. Details of each cell line used. Insertion and unedited allele(s) were detected using PCR and genomic sequencing. KSW, Knocksideways.

| Cell line | Clone | Genomic status | Western band-shift | Western endogenous | Insertion | Unedited allele(s) | Expected localization | KSW | Mitosis | Notes |
| --- | --- | --- | --- | --- | --- | --- | --- | --- | --- | --- |
| CLTA-FKBP-GFP | 5 | Homozygous | Yes | No | Yes | No | Yes | Yes | Normal | Clathrin triskelia are CLTC with CLTA and CLTB. In HeLa, CLTA > CLTB expression |
| GFP-FKBP-TACC3 | D5 | Homozygous | Yes | No | Yes | No | Yes | Yes | Slight delay | – |
| chTOG-FKBP-GFP | H5 | Heterozygous | Yes | Yes | Yes | Yes | Yes | Yes | Normal | >20 heterozygous clones, 0 homozygous recovered from three attempts |
| GTSE1-FKBP-GFP | B5 | Homozygous | Yes | No | Yes | No | Yes | Yes | Normal | – |

**Table 2.** Summary of knocksideways experiments. Each row is a cell line and the effect of relocalization of the tegged protein to mitochondria on the spindle localization of each protein-of-interest is indicated.

| Cell line | Protein of interest |  |  |  |  |  |  |  |  |  |  |  |
| --- | --- | --- | --- | --- | --- | --- | --- | --- | --- | --- | --- | --- |
|  | clathrin |  |  | TACC3 |  |  | chTOG |  |  | GTSE1 |  |  |
|  | IF | Live | Conclusion | IF | Live | Conclusion | IF | Live | Conclusion | IF | Live | Conclusion |
| CLTA-FKBP-GFP | X | X | X | ↓↓ | ↓↓ | ↓↓ | – | ↓↓ | ↓↓ | ↓ | ↓↓ | ↓↓ |
| GFP-FKBP-TACC3 | ↓ | ↓ | ↓ | X | X | X | – | ↓↓ | ↓↓ | ↓ | ↓↓ | ↓↓ |
| chTOG-FKBP-GFP | – | – | – | – | – | – | X | X | X | – | – | – |
| GTSE1-FKBP-GFP | – | – | – | – | – | – | – | – | – | X | X | X |

**Figure S1.**
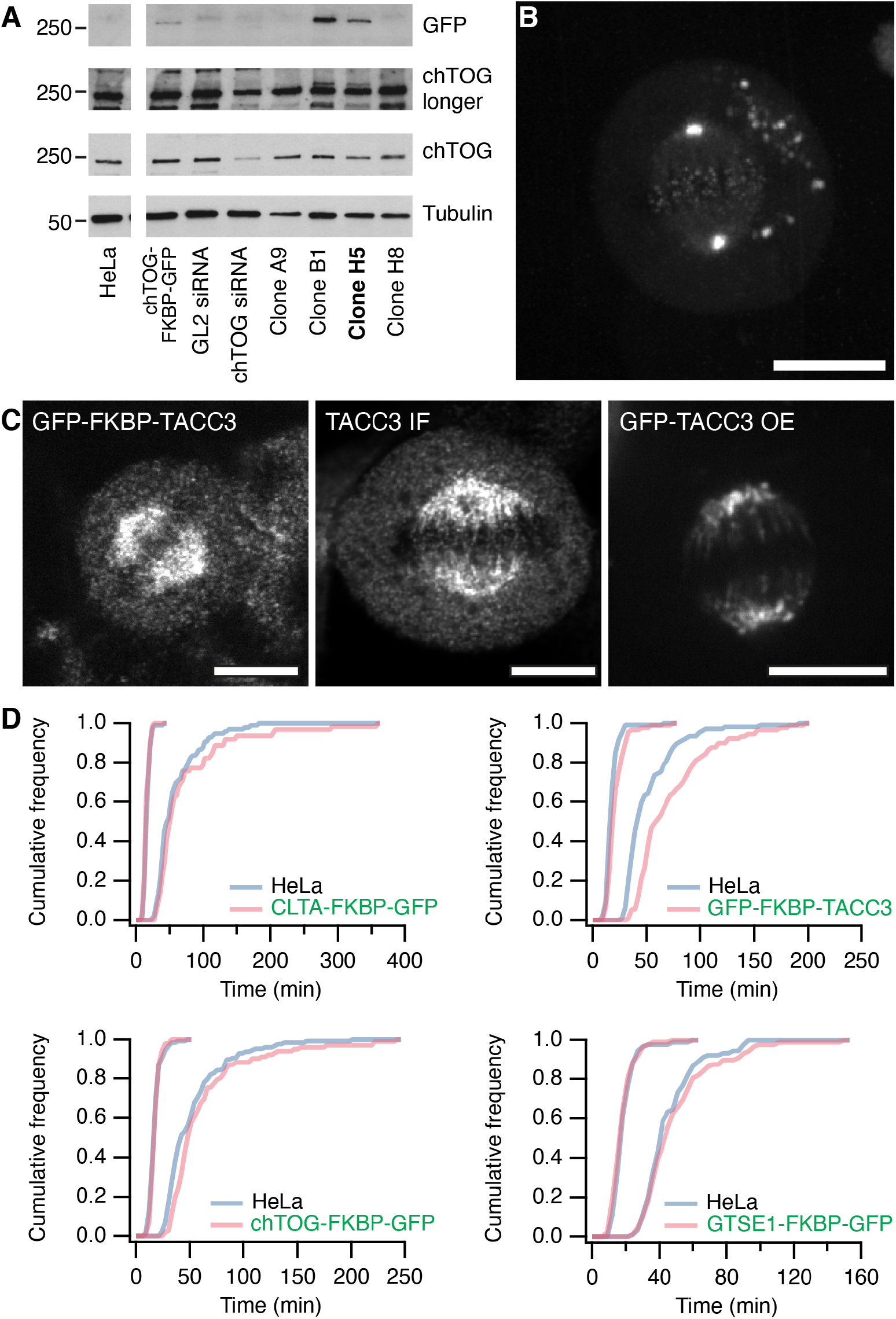
Further validation of gene edited cell lines. (**A**) Western blots of four representative chTOG-FKBP-GFP clones. For comparison, parental HeLa are shown, untransfected (HeLa) or expressing chTOG-FKBP-GFP, or transfected with siRNA as described. A single gel is shown probed with a chTOG antibody (two different exposures), tubulin as a loading control, the upper blot was reprobed with a GFP antibody. The chTOG antibody does not appear to detect the tagged protein with the same efficiency as the unedited protein. (**B**) Maximum intensity projection of a stack of confocal images of a live chTOG-FKBP-GFP knock-in cell at metaphase. (**C**) Representative confocal micrographs of the GFP-FKBP-TACC3 cell line (with GFP booster), anti-TACC3 immunofluorescence in unedited HeLa cells, and over-exprsesion of GFP-TACC3 in unedited HeLa. Scale bar, 10 μm. (**D**) Mitotic progression of gene-edited cell lines. Cumulative histograms of timings from nuclear envelope breakdown to metaphase (NEB-M, long duration) and metaphase to anaphase (M-A, short duration). Gene-edited cells are as indicated and were imaged alongside their respective unedited parental HeLa counterpart. All imaging experiments were done three times. Number of cells analyzed (edited line and parental) = CLTA-FKBP-GFP: 62 and 97; GFP-FKBP-TACC3: 90 and 106; chTOG-FKBP-GFP: 102 and 128; GTSE1-FKBP-GFP: 88 and 90.

**Figure S2.**
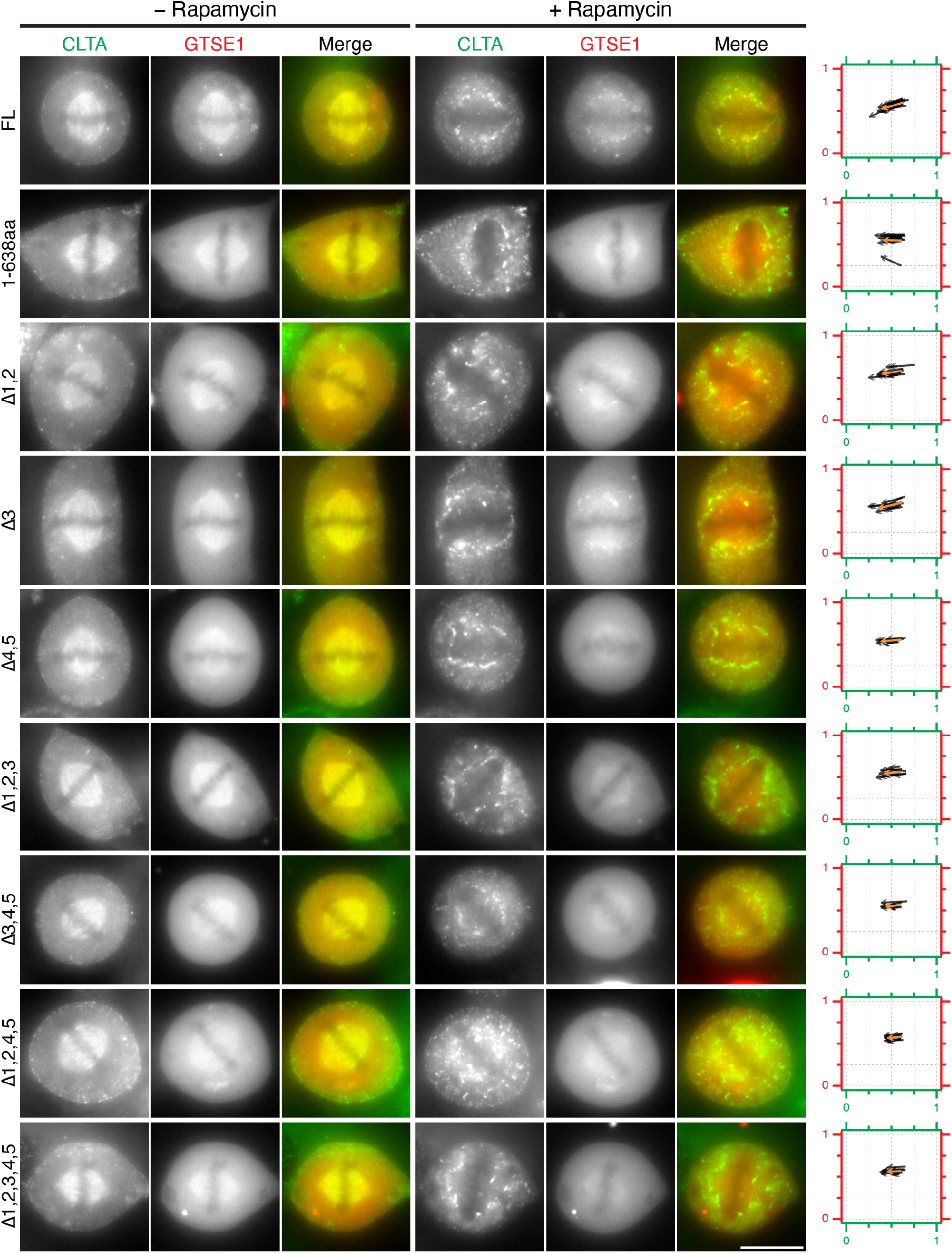
Live cell imaging of knocksideways in CLTA-FKBP-GFP cells expressing GTSE1 LIDL mutants. Stills from live cell imaging of clathrin knocksideways in metaphase CLTA-FKBP-GFP cells expressing the indicated GTSE1-mCherry constructs. Rapamycin (200 nM) was added to induce removal of clathrin and imaged for a total of 10 min to visualize co-rerouting of GTSE1 mutants. Scale bar, 10 μm. (Right) Quantification of GTSE1 co-rerouting shown as arrow plots. Arrows show the fraction of spindle and mitochondria fluorescence that is at the spindle (i.e. 1 = completely spindle-localized, 0 = mitochondria-localized), for both channels, moving from pre to post rapamycin localization. Black arrows represent individual cells, the orange arrow is the mean. n = 7-12 cells per condition.

**Figure S3.**
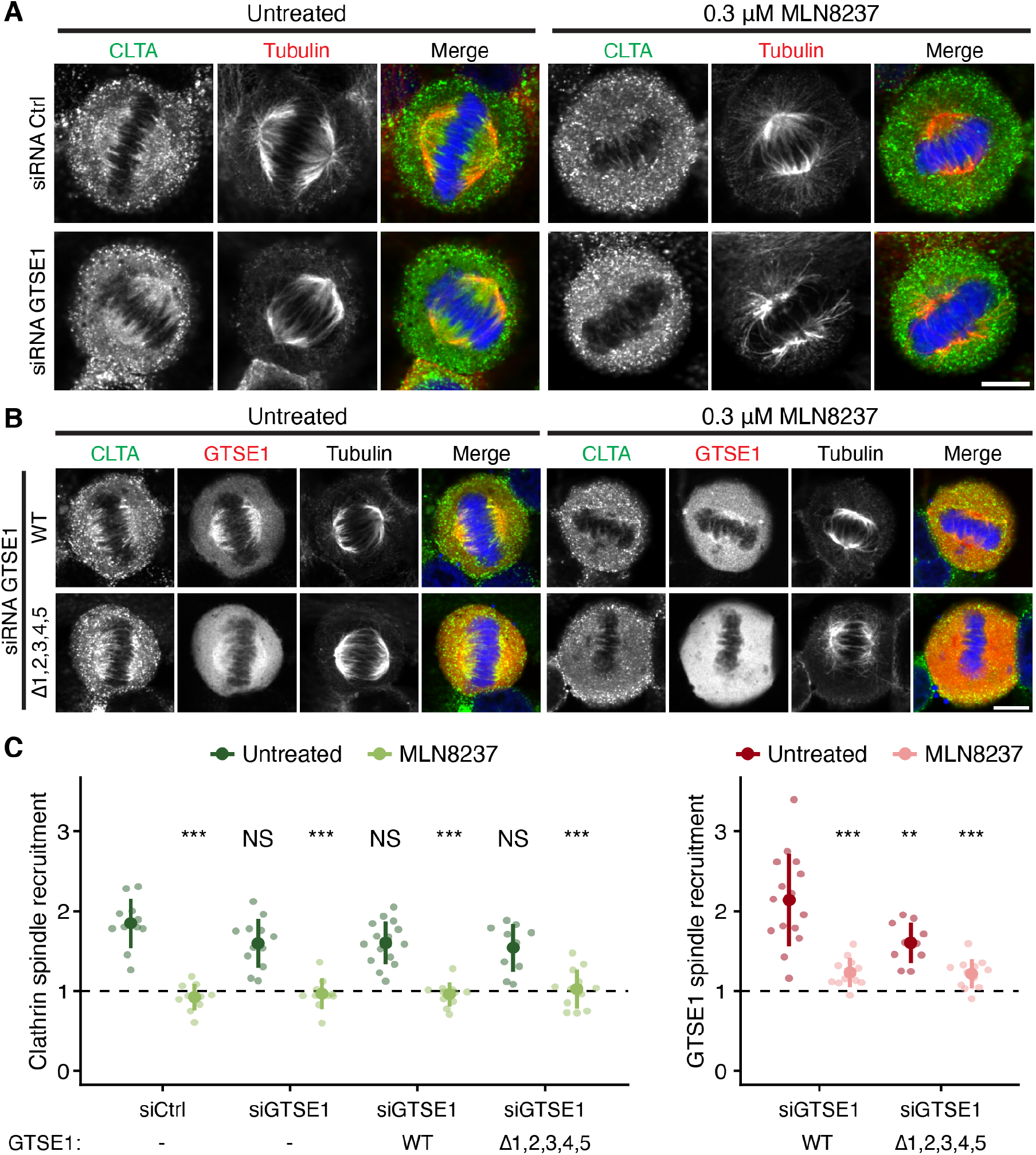
Comparison of GTSE1 LIDL motif ablation with the effect of Aurora-A inhibition on spindle localization of clathrin and GTSE1. Representative widefield micrographs of CLTA-FKBP-GFP cells at metaphase to show the spindle localization of clathrin (**A**), or GTSE1-mCherry construct (WT or Δ1,2,3,4,5, red) and clathrin (**B**). Cells were treated with control (GL2, Ctrl) or GTSE1 siRNA and Aurora-A kinase was inhibited with MLN8237 (0.3 μM, 40 min) as indicated. Cells were stained for tubulin (red in A, not shown in merge in B) and DNA (blue). A GFP-boost antibody was used to enhance the signal of CLTA-FKBP-GFP (green). Scale bar, 10 μm. (**C**) Quantification of clathrin and GTSE1 spindle recruitment. Each dot represents a single cell, n = 10-15 cells per condition. The large dot and error bars show the mean and the mean ±SD, respectively. Analysis of variance (ANOVA) with Tukey’s post-hoc test was used to compare the means between each group, using the untreated cells + siRNA Ctrl (clathrin) and untreated cells + WT GTSE1 (GTSE1) for comparison. The p-value level is shown compared to WT: ***, *p* < 0.001; **, *p* < 0.01; NS, *p* > 0.05.

**Figure S4.**
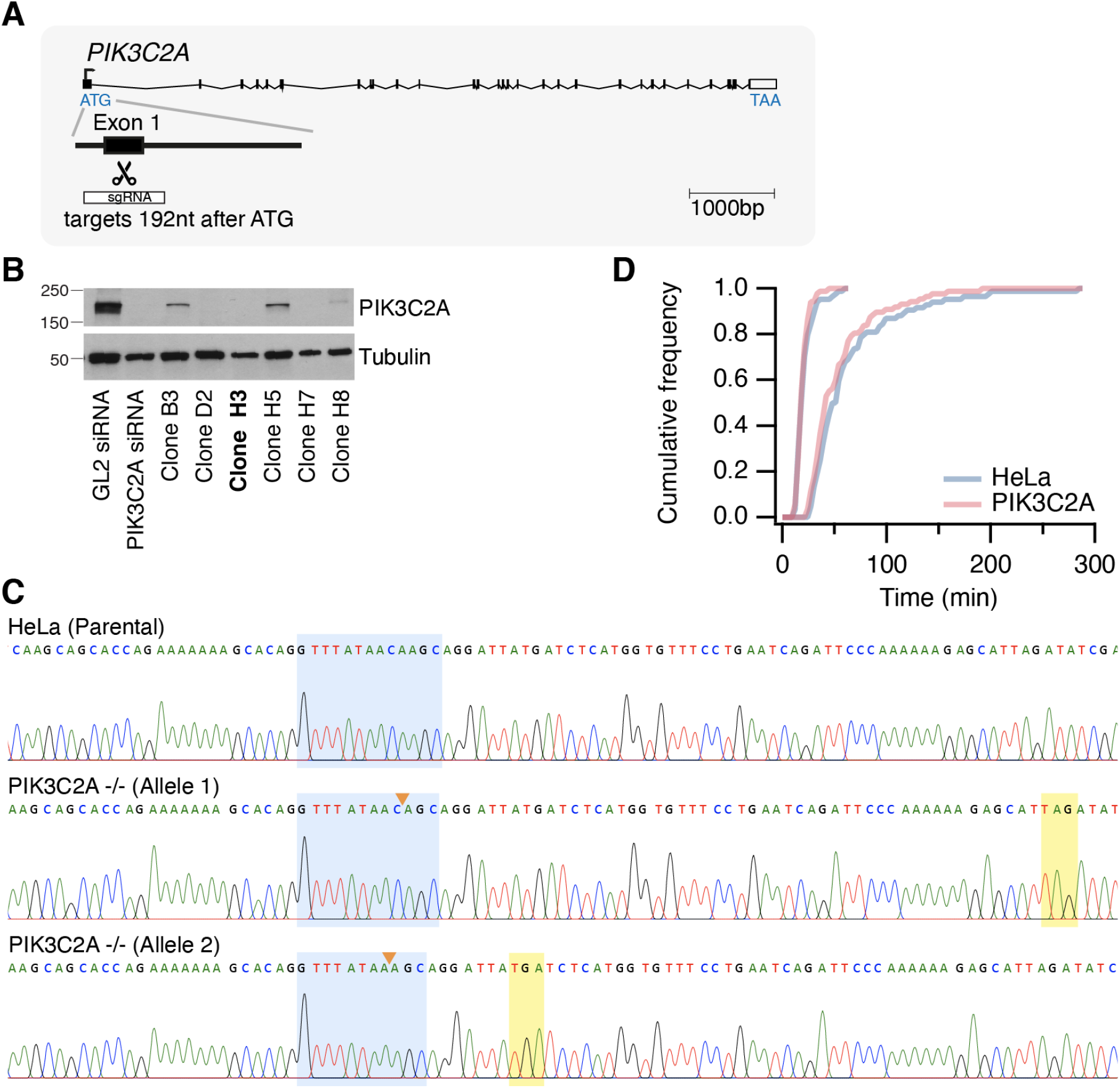
Generation of PIK3C2A-null HeLa cells. (**A**) Targeting strategy for generation of a PIK3C2A-null cell line. HeLa cells were transfected with plasmid to express GFP coupled Cas9 nuclease and sgRNA targeting 192 bp from the start codon of the PIK3C2A gene. Scale bar, 1000 bp. (**B**) Western blot of a selection of clones grown after expansion of GFP-expressing cells sorted by FACS. Presence of a band for PIK3C2A was assessed compared parental HeLa cells treated with PIK3C2A siRNA or control, GL2. Tubulin, loading control. Clone H3, was used in this study (bold). (**C**) A genomic fragment from clone H3 was cloned into a cloning vector and 20 bacterial clones were picked and sequenced to assess the status of PIK3C2A alleles. We found two sequences, indicating two alleles and both had deletions (orange arrows) which resulted in premature truncation of the PI3KC2A gene after 87 and 72 residues, respectively. Stop codon is highlighted in yellow. Blue window highlights the edited region. (**D**) Mitotic progression of PIK3C2A-null cells compared to parental HeLa cells. Cumulative histograms of timings from nuclear envelope breakdown-to-metaphase (NEB-M, long duration) and metaphase-to-anaphase (M-A, short duration). Progression experiments were done three times. Number of cells analyzed = PIK3C2A^−/−^: 87; parental: 84.

**Figure S5.**
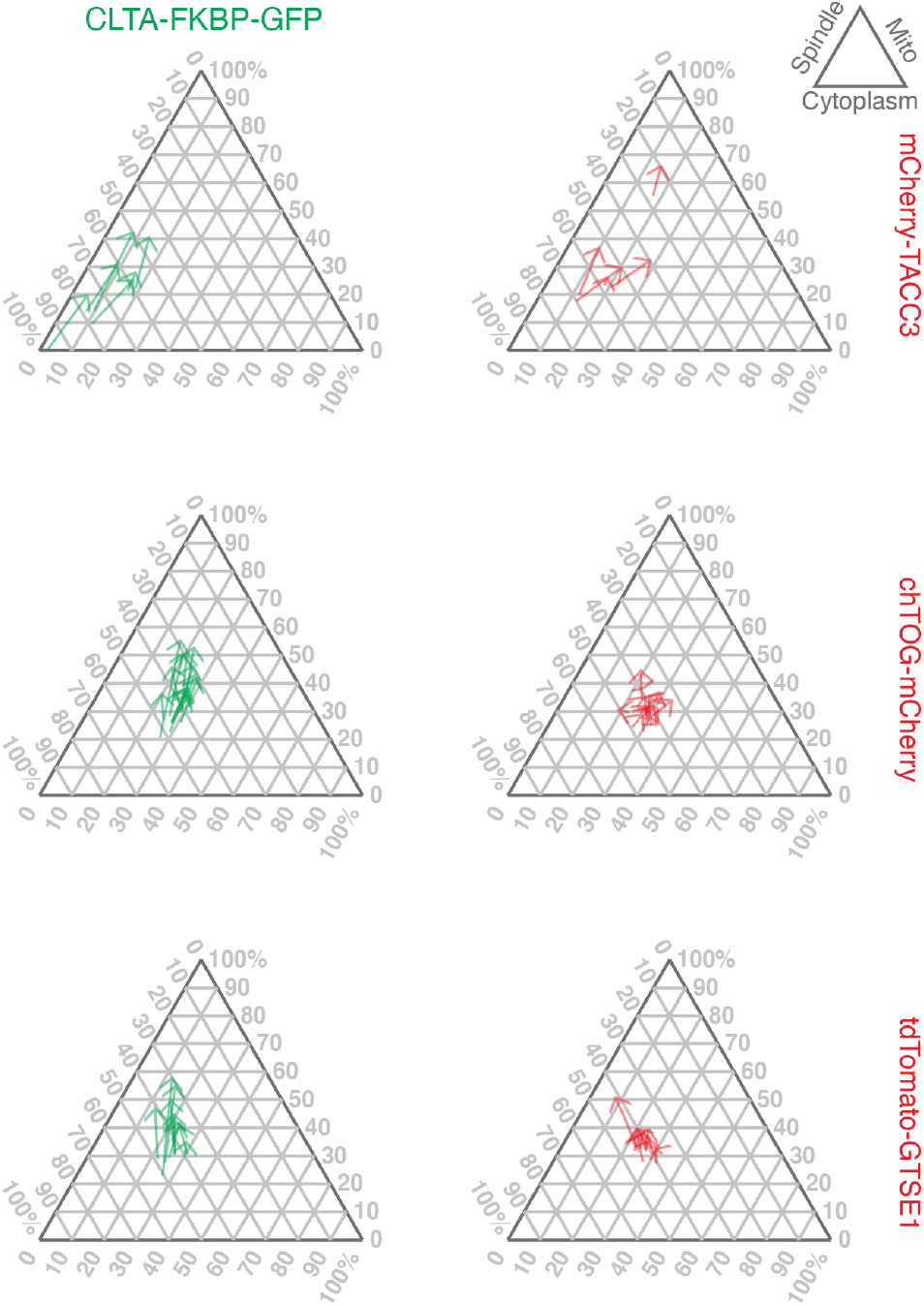
Ternary diagrams of live CLTA-FKBP-GFP spatial relocalization experiments. Localization before and after addition of rapamycin is shown by an arrow for each cell. Ternary diagrams can be read using the key. For example, a protein that is localized entirely on the miochondria and is absent from the spindle would be at the top corner of the triangle. Generally, movement (if it occurs) is from the bottom left corner to the upper corner with cytoplasmic signal staying approximately constant.

### Supplementary Videos

**Figure SV1.**
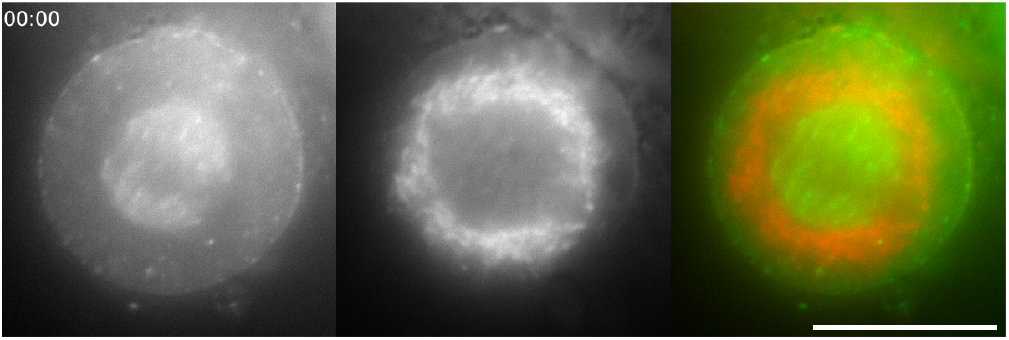
Knocksideways of CLTA-FKBP-GFP. Typical widefield movie of relocalization in response to rapamycin (200 nM) in cells co-expressing mCherry-MitoTrap. Time, mm:ss. Scale bar, 10 μm.

**Figure SV2.**
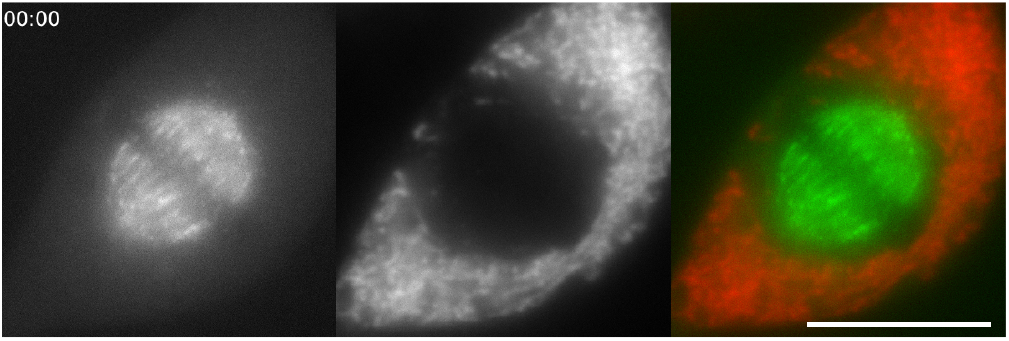
Knocksideways of GFP-FKBP-TACC3. Typical widefield movie of relocalization in response to rapamycin (200 nM) in cells co-expressing mCherry-MitoTrap. Time, mm:ss. Scale bar, 10 μm.

**Figure SV3.**
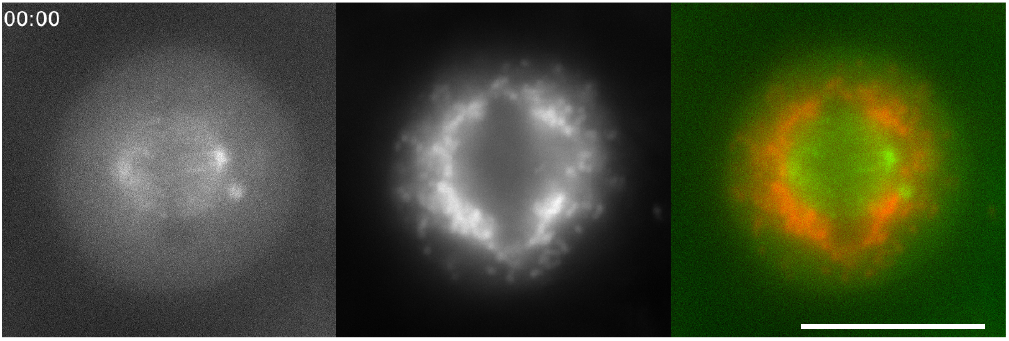
Knocksideways of chTOG-FKBP-GFP. Typical widefield movie of relocalization in response to rapamycin (200 nM) in cells co-expressing mCherry-MitoTrap. Time, mm:ss. Scale bar, 10 μm.

**Figure SV4.**
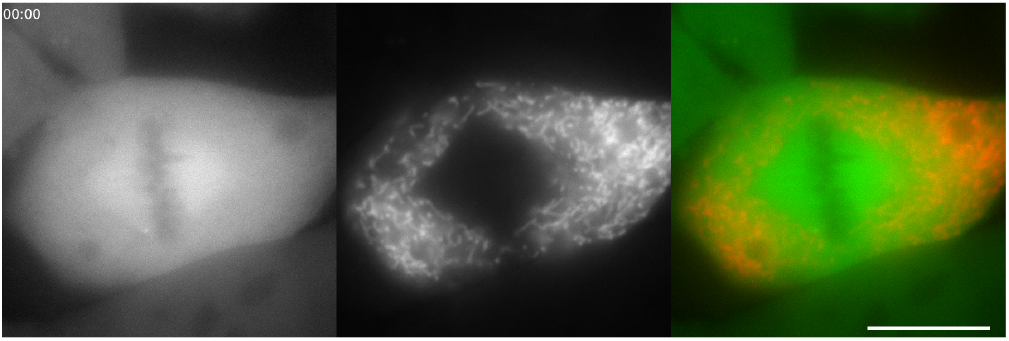
Knocksideways of GTSE1-FKBP-GFP. Typical widefield movie of relocalization in response to rapamycin (200 nM) in cells co-expressing mCherry-MitoTrap. Time, mm:ss. Scale bar, 10 μm.

